# Neuroprotective Effect of Combined Pomegranate and Candesartan Therapy Against Chronic Cerebral Ischemia in Rats

**DOI:** 10.64898/2026.02.23.707366

**Authors:** Rana Awada, Fatima Radi, Zaher Abdel Baki, Akram Hijazi, Wissam H. Joumaa, Zeinab Ezzeddine, Laurent. O. Martinez, Mohamad Nasser

**Author notes:** **Corresponding Authors:** (Mohamad Nasser) (Laurent Martinez).

## Abstract

**Background:** Stroke remains a leading cause of mortality worldwide. Ischemic stroke, caused by arterial occlusion, induces sensorimotor deficits and memory impairments. Excessive activity of the brain angiotensin II type 1 receptor (AT1R) is associated with hypertension and cerebral ischemia. Candesartan (CN), an AT1R blocker, improves cerebrovascular blood flow. Pomegranate (*Punica granatum*) is rich in polyphenolic antioxidants that reduce oxidative stress and inflammation, suggesting potential neuroprotective effects in cerebral ischemia.

**Aim:** This study compared the neuroprotective effects of CN administered alone or in combination with pomegranate (POM) in a rat model of cerebral ischemia induced by chronic unilateral carotid artery ligation.

**Methods:** Cerebral ischemia was induced by ligation of the right common carotid artery (RCCA) in adult rats. Animals were randomly assigned to four groups: sham control, untreated ischemic, ischemic treated with CN, and ischemic treated with CN + POM. Sensorimotor and cognitive functions were assessed 1–15 days post-surgery using beam balance (BB), beam walking (BW), modified sticky-tape (MST), novel object recognition (NOR), and the Morris water maze (MWM) tests.

**Results:** RCCA ligation induced marked sensorimotor deficits and memory impairments. Both CN monotherapy and CN + POM treatment equally restored sensorimotor function to sham-control levels, as demonstrated by BB, BW, and MST tests. In contrast, CN + POM treatment showed greater efficacy than CN alone in improving short-term recognition and spatial memory, as demonstrated by NOR and MWM performance.

**Conclusion:** CN effectively reverses ischemia-induced sensorimotor deficits, whereas the addition of POM confers specific and enhanced protection against cognitive impairment, indicating distinct mechanisms underlying sensorimotor and memory recovery after cerebral ischemia.

## 1. Introduction

Stroke is characterized by a sudden disruption of blood supply to the brain, resulting from either hemorrhagic or ischemic events, which leads to severe neurological deficits [1]. Its global incidence remains high and is projected to increase significantly over the coming decades [2]. Ischemic stroke accounts for approximately 80% of all cases, frequently resulting in long-term impairment of physical and cognitive functions, including deficits in learning and memory [3].

The renin-angiotensin system (RAS) is a crucial hormonal pathway and a primary regulator of blood pressure, cardiovascular system, and neurodegenerative processes [4]. In animal models of focal cerebral ischemia, RAS modulation has been shown to improve neurological outcomes and reduce brain injuries by attenuating the signaling pathway of the angiotensin II type 1 receptor (AT1R), the main effector of this system [4]. Candesartan (CN), an AT1R antagonist, has demonstrated significant neuroprotective efficacy in stroke animal models. CN improves neurobehavioral outcomes by reducing infarct size [5]. When administered intravenously post-ischemia, CN has been shown to enhance vascular remodeling and provide sustained functional benefits [6].

Beyond hemodynamic regulation, the pathophysiology of cerebral damage is driven by the production of reactive oxygen species (ROS) and other toxic metabolite produced by the brain during ischemia [7]. While the brain possesses an endogenous antioxidant defense system to neutralized these free radicals, is often overwhelmed during chronic ischemia [8]. Consequently, there is a growing interest in plant-derived bioactive compounds for their therapeutic potential [9]. Natural extracts can safeguard neurons from ischemic damage through their antioxidant properties [10]. Specifically, polyphenols have emerged as effective neuroprotectors in various models of cerebral ischemia and periventricular white matter lesions [11]. Resveratrol, a polyphenol found in various plants, including grapes, plums and peanuts has shown various medicinal properties, including antioxidant, protection of cardiovascular disease and cancer risk [12]

This study aims to investigate the neuroprotective effects of combining pomegranate juice with candesartan against cerebral ischemia induced by chronic unilateral common carotid artery ligation in rat. Furthermore, we seek to determine whether pomegranate juice can potentiate the therapeutic efficacy of candesartan in restoring cognitive and sensorimotor functions.

## 2. Materials and Methods

### 2.1. Experimental animals and grouping

Twenty-four adult male Wistar rats (280-320 g) were randomly assigned to four experimental groups (n = 6 per group): (i) a sham-operated control group subjected to surgical incision without ligation; (ii) an untreated ischemic group subjected to right common carotid artery (RCCA) ligation; (iii) a candesartan (CN) group treated with candesartan (0.5 mg/kg/day); and (iv) a combination treatment (CN + POM) group receiving candesartan (5 mg/kg/day) and pomegranate juice extract (400 mg/kg/day). Sham-operated and untreated ischemic groups received drinking water without treatment. All animals were housed under controlled environmental conditions (a 12-h light/dark cycle, temperature 23 ± 1°C) with *ad libitum* access to standard chow and water. Rats were euthanized on post-operative day 16.

### 2.2. Induction of chronic unilateral ischemia

Rats were anesthetized via intraperitoneal injection of a mixture of ketamine (100 mg/kg) and xylazine (3 mg/kg) and placed on a surgical table in a supine position. Following cervical shaving and disinfection, a 25-mm midline incision was made from the mandible base to the sternum. The RCCA was carefully isolated from the surrounding nerves and soft tissues. The artery was then occluded using 4-0 silk sutures with two firm knots to ensure complete ligation without damaging the arterial wall. The surgical site was then closed with nylon sutures. Sham animals underwent the same surgical exposure of the RCCA without ligation.

### 2.3. Treatment preparation and administration

#### 2.3.1. Pomegranate extract

Pomegranates (*Punica granatum L.*) were sourced from Nabatieh, Lebanon (approximately 400 m altitude). The fruits were pressed to obtain juice, which was then filtrated, and stored at −80°C for 2–3 hours. The juice was then lyophilized for 72 hours to produce a fine powder. This extract was stored in desiccators at room temperature until its use for chemical analysis and administration.

#### 2.3.2. Pharmacological treatment

Candesartan cilexetil (Atacand, Takeda Pharmaceutical) was administered at a dose of 0.5 mg/ kg/ day via drinking water [6]. In the combination group, pomegranate extract was administrated at a dose of 500 mg/ kg/ day via drinking water, based on established protocols [13].

### 2.4. Chemical analyses

#### 2.4.1. Photochemical screening tests

Pomegranate juice was centrifuged at 2,500 rpm (22°C) and filtered using a Bushner aspirator. The filtrate was analyzed for primary and secondary metabolites, including flavonoids and alkaloids, using standard qualitative screening methods, as detailed in [14] and presented in **Tables S1 and S2** of the Supplementary Material.

#### 2.4.2. DPPH radical scavenging assay

The antioxidant capacity of POM extract was measured using the *2,2-Diphenyl-1-picrylhydrazyl* (DPPH) assay. Two concentrations of the extract (10 mg/mL and 20 mg/mL) were prepared. Extracts were dissolved in a mixture of water and methanol (0.5 mL each). For the assay, 50 µL of each extract was mixed with 2 mL of DPPH solution (0.0012 g in 50 mL of methanol). The reaction mixtures were vortexed and incubated for 30 min at room temperature in the dark. Absorbance was measured at 515 nm using a spectrophotometer (UV–Vis. Cary 4000 UV-Vis, Agilent, UK). Ascorbic acid (ABS) was used as a positive control. The control solution consisted of 2 mL DPPH solution mixed with solvent only (without extract).

Radical scavenging activity was calculated according to the following equation:

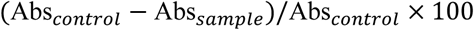

where Abs *_control_*is the absorbance of the DPPH solution with solvent, and Abs*_sample_*is the absorbance of the DPPH solution in the presence of the extract.

#### 2.4.3. Total carbohydrate content determination

A total of 100 mg of pomegranate powder was macerated with 5.25 mL of 80% ethanol for 12 h, then centrifuged at 4000 rpm for 10 minutes. The supernatant containing the sugars was collected and diluted 30-fold. A tenth of this diluted solution was further diluted in water to prepare Solution A. Solution B was prepared by dissolving 1 g of anthrone in 500 ml of concentrated sulfuric acid. For the assay, 2 mL of Solution A was mixed with 4 mL of solution B in a tube, vortexed for one minute and then incubated at 92°C for 10 minutes. After incubation, the tubes were cooled and kept in the dark for 30 minutes. Absorbance was measured at 585 nm.

#### 2.4.4. Total phenolic content (TPC)

TPC was determined using the Folin-Ciocalteau method [15]. Briefly, 25 mg of pomegranate powder was dissolved in 10 mL of distilled water. An aliquot of 40 µL of the extract was mixed with 3.16 mL of distilled water and 200 µL of Folin-Ciocalteau reagent. After 5 minutes, 400 µL of sodium carbonate solution (7.5% w/v) was added, and the mixture was incubated at 40°C for 30 minutes. Absorbance was then measured at 765 nm against a blank.

TPC was calculated using the gallinc acid calibration curve and expressed as galling acid equivalent (GAE):

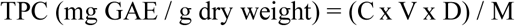

where C is the concentration of gallic acid (mg/mL) determined form the calibration curve, V is the volume of the extract (mL), D is the dilution factor, and M is the mass of the sample (g, dry extract).

### 2.5. Behavioral and memory Assessments

#### 2.5.1. Beam Balance (BB) test

The BB test was employed to assess static equilibrium, postural control and vestibulomotor function. Rats were trained to balance on a wooden beam (66 cm in length, 1.5 cm in diameter, elevated 70 cm above the floor) for four days, three times per day, prior to surgery. The maximum duration of each training session was 60 seconds. Following surgery, performance was assessed by conducting the test three times per testing day, considered as technical triplicates. Performance was scored according to the following scale: stable balance (1), shaky balance (2), attempting to balance by hanging with 1 or 2 limbs (3), trying to balance but falling after 10 seconds (4), falling off in under 10 seconds (5) and falling off without any effort to balance or hang onto the beam (6) [16]. The mean score of technical triplicates was used for analysis.

#### 2.5.2. Beam walking (BW) test

Sensorimotor coordination and dynamic balance were evaluated using the BW test. The apparatus consisted of a wooden beam (156 cm long, 1.6 cm wide, 3 cm high) supported by two stands and elevated 100 cm above the ground. A starting wooden platform (30 cm x 30 cm) was positioned at one end of the beam, while the opposite end led to a darkened escape box (30 cm x 30 cm x 13.5cm). Prior to surgery, rats underwent a standardized habituation protocol. Animal were first placed inside the escape box for 5 minutes, followed by placement on the beam at progressively increasing distances from the box to encourage forward locomotion. Finally, rats were placed on the starting platform to traverse the full length of the beam [17]. Training was conducted daily until animals achieved a criterion of optimal performance, defined as a crossing latency of 4–6 seconds. The test was repeated on post-operative days +1, +3, +5, and +7. During each session, three trials were performed per rat, and crossing time was recorded. The results were averaged for analysis.

#### 2.5.3. Modified sticky tape (MST) test

The MST test was used to assess forelimb somatosensory function and sensorimotor integration on the ipsilateral and contralateral sides. A piece of sticky tape measuring 5–6 cm in length and 1.3 cm in width was wrapped around the forepaw of the rat. Two timers were used simultaneously: the first ran continuously for 30 seconds, while the second was turned on when the rat attempted to remove the tape during this period [18]. The test was conducted in the rat’s home cage under quiet conditions. Training was performed for three days prior to surgery, with three trials per forelimb each day. On the final day of training, all rats were required to detect the tape within 30 seconds for both forelimbs. The test was conducted after the BB and BW tests on postoperative days +1, +3, +5, and +7.

#### 2.5.4. Novel object recognition (NOR) test

The NOR test was used to assess recognition memory and was conducted on postoperative day 8. Initially, each rat was placed in a black box (51.5 cm x 40.5 cm x 23.5 cm) for 5 minutes for habituation. Two identical objects (Duplo® Toys) were then placed equidistant from the rat, and the time spent exploring each object was recorded over a 5-minute period. The rat was subsequently returned to its home cage for 5 minutes before being reintroduced into the box, where one of the familiar objects was replaced with a novel object of a different shape. Exploring times for both the familiar and novel objects was recorded during a second 5-minute session [19]. This test, lasting a total of 20 minutes per rat, was used to evaluate short-term memory.

#### 2.5.5. Morris water maze (MWM) test

Long-term spatial memory was assessed using the MWM. The apparatus consisted of a circular pool (183 cm in diameter, 38 cm in height) filled with water mixed with milk to a depth of 21 cm. A hidden platform (13 cm in diameter, 20 cm in height) was submerged 1 cm bellow the water surface and positioned 17–18 cm from the pool wall. Visual cues were placed around the pool to help spatial navigation. The pool was virtually divided into four quadrants. On day 1, the platform was placed in one quadrant, and its position was changed to a different quadrant on each subsequent day. Each trial began by placing the rat on the platform for 20 seconds, followed by release into the pool from the quadrant opposite the platform. The latency to locate the platform was recorded. If the rat failed to find the platform within 60 seconds, it was gently guided to it and allowed to remain there for 20 seconds. The rat was then removed and dried under a warm heat source. Each rat underwent three trials per day, starting from three different quadrants, for seven consecutive days (postoperative days +10 to +15) [20].

### 2.6. Statistical analyses

Statistical analyses were performed using GraphPad Prism V.10.6.1 (GraphPad Software). Data are presented as mean ± SEM for the indicated number of observations. Prior to statistical testing, data normality was assessed using the Shapiro-Wilk test. As all datasets were normally distributed, differences between groups were evaluated using a one-way ANOVA followed by Tukey’s post hoc test for multiple comparisons. A P-values < 0.05 was considered statistically significant.

### 2.7 Ethical Approval

Animal experiments were conducted humanely in accordance with the Regulations for Animal Experiments of Islamic University of Lebanon, which comply with the revised Animals (Scientific Procedures) Act 1986 (UK) and Directive 2010/63/EU (Europe). The study protocol was approved by the Research Ethics Committee of the Islamic University of Lebanon (reference number: IUL-EC-25-A005).

## 3. Results

### 3.1. Phytochemical screening and antioxidant activity of pomegranate (POM) fruit extract

Phytochemical screening was conducted on extracts obtained from the seeds and juice of Lebanese pomegranates cultivated at approximately 400 m above sea level. As shown in **Table S2** of the Supplementary Material, both seed-and juice-derived POM extracts contained flavanones, flavonoids, terpenoids, lignins, alkaloids, phenols, sterols, and steroids. Reducing sugars, resins, quinones, and cardiac glycosides were found exclusively in the seed extract, whereas tannins and anthocyanins were identified only in the juice extract. Based on its higher content of bioactive compounds and superior antioxidant activity, the juice-derived extract was selected for subsequent experiments.

The antioxidant capacity of the juice extract was assessed using the DPPH free radical scavenging assay. The scavenging activity increased in a dose-dependent manner relative to POM concentration (**Table 2**). Further analysis confirmed that the POM juice extract is highly enriched in carbohydrates and phenolic compounds (**Table 3**).

**Table 1.** Dose-dependent antioxidant capacity of pomegranate juice extract as determined by the DPPH free radical scavenging assay.

|  | **Control**<br>(DPPH and water) | **POM juice**<br>(10 mg/ml) | **POM juice**<br>(20 mg/ml) |
| --- | --- | --- | --- |
| **Scavenging Activity (%)** | 0 | 28 | 55 |

*Abbreviations: DPPH, 2,2-Diphenyl-1-picrylhydrazyl; POM, pomegranate.*

**Table 2.** Carbohydrates and phenol content in the pomegranate juice extract.

| **Test** | **Content** |
| --- | --- |
| **Carbohydrates (%)** | 69.3 ± 2.5 |
| **Total phenol (mg GAE/g DW)** | 7 ± 0.09 |

*Values are expressed as % or mean ± SEM.*
*Abbreviations: DW, dry weight; GAE, gallic acid equivalence.*

### 3.2. Beam Balance (BB) test

The BB test was used to evaluate static balance and postural stability. Baseline performance was established on day-1 (final training day), when all rats achieved the optimal score of 1 (**Figure 1**).

**Figure 1.**
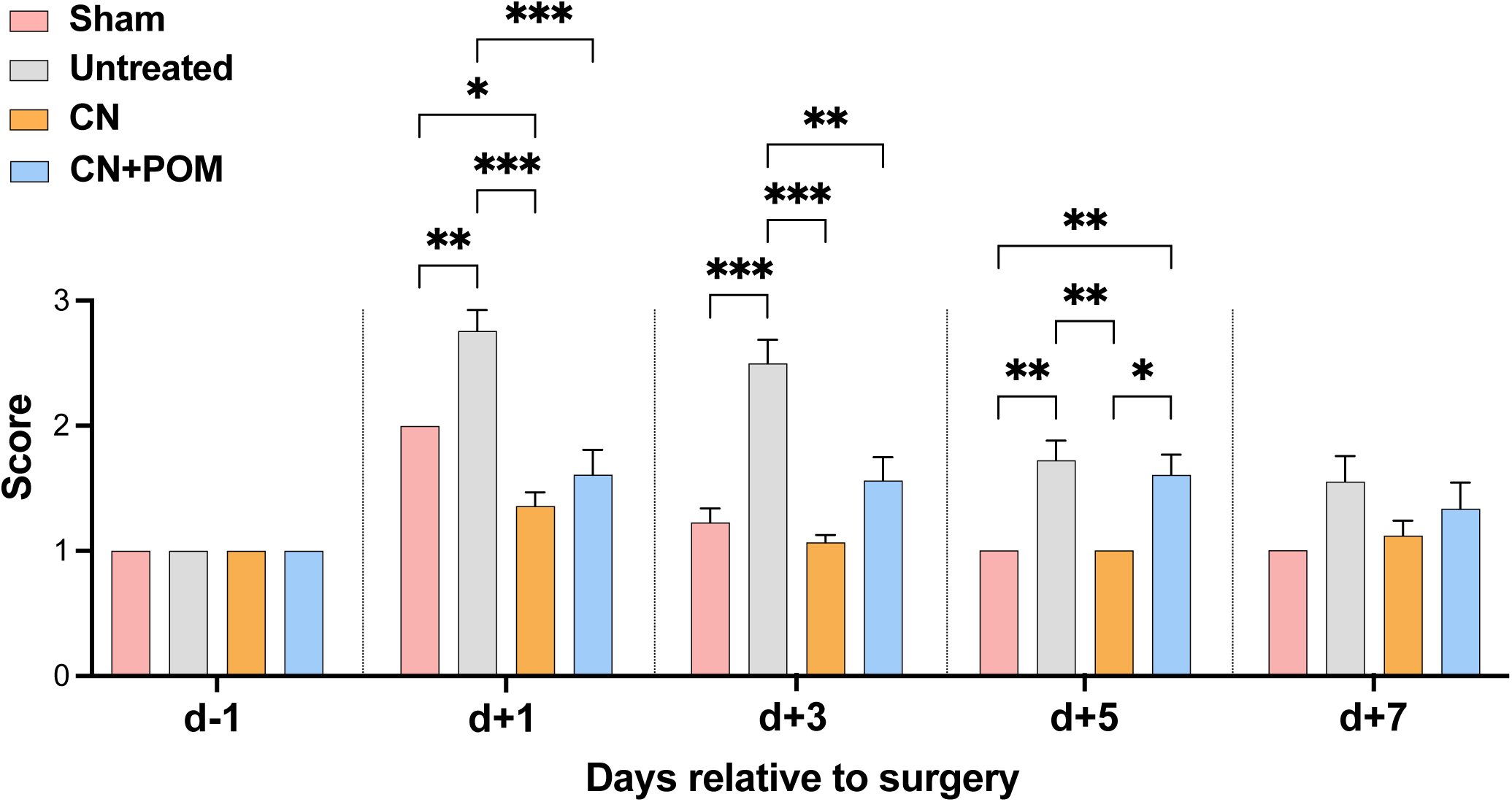
Beam Balance (BB) test. Beam Balance scores were measured at baseline (day −1) and on post-operative days +1, +3, +5, and +7 in all experimental groups. Data are presented as mean ± SEM. Statistical comparisons were performed independently at each time point using one-way ANOVA followed by Tukey’s post hoc test (*P < 0.05, **P < 0.01, ***P < 0.001). Group and sample size: Sham, sham-operated control group subjected to surgical incision without ligation (n = 6); Untreated, untreated ischemic group subjected to RCCA ligation (n = 6); CN, CN-treated ischemic group (n = 5); CN+POM, ischemic group treated with the combination of CN and POM (n = 6). Abbreviation: RCCA, right common carotid artery; CN, candesartan, POM, pomegranate.

From post-operative day +1 to day +3, the untreated ischemic group showed a significant deficit compared to the sham control group (P < 0.001 and P < 0.01, respectively). This deficit was followed by spontaneous and progressive functional recovery, with performance approaching sham levels by days +5 and +7.

Both CN monotherapy and the combined CN + POM treatment significantly restored BB performance compared with the untreated ischemic group, with a significant effect already evident on day +1. No additional synergistic benefit was observed with the combined treatment compared with CN alone. The treatment effects were most pronounced on days +1 and +3, while performance converged across all experimental groups by days +5 and +7, consistent with spontaneous recovery in untreated animals.

### 3.3. Beam Walking (BW) test

The BW test assesses sensorimotor coordination and dynamic balance during locomotion (**Figure 2**).

**Figure 2.**
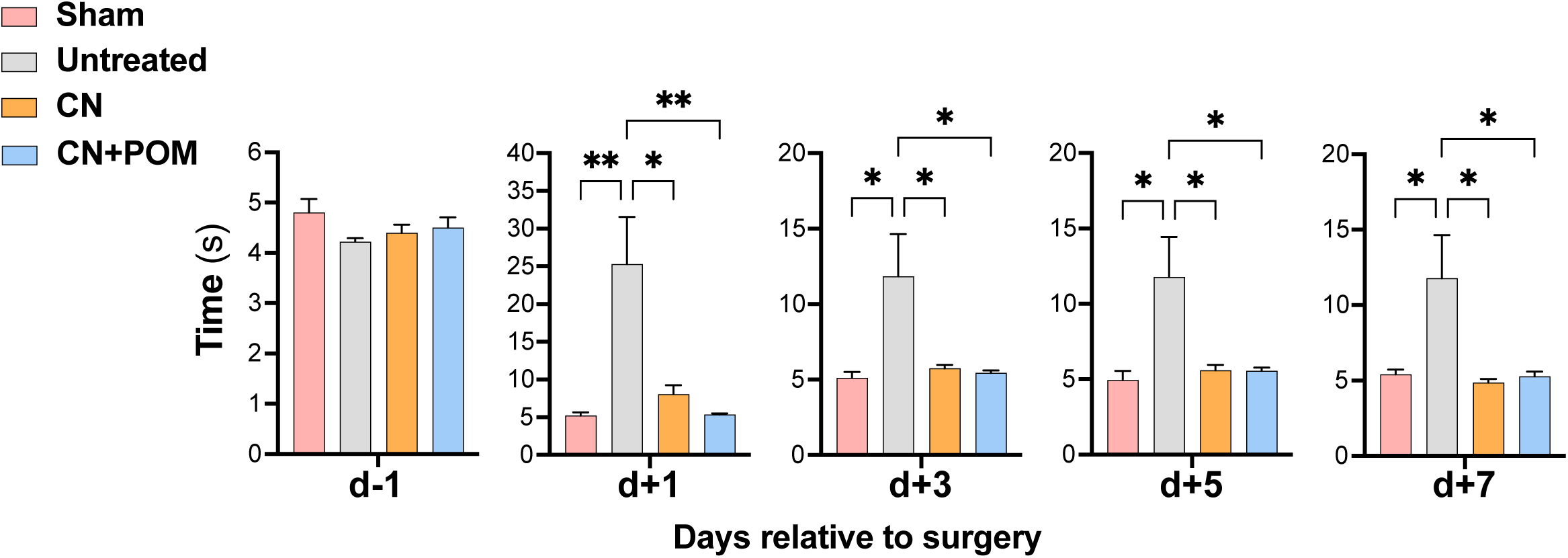
Beam Walking (BW) test. Beam Walking crossing latency was measured at baseline (day −1) and on post-operative days +1, +3, +5, and +7 in all experimental groups. Data are presented as mean ± SEM. Statistical comparisons were performed independently at each time point using one-way ANOVA followed by Tukey’s post hoc test (*P < 0.05, **P < 0.01). Group and sample size: Sham, sham-operated control group subjected to surgical incision without ligation (n = 6); Untreated, untreated ischemic group (n = 6); CN, CN-treated ischemic group (n = 5); CN+POM, ischemic group treated with the combination of CN and POM (n = 6). Abbreviation: CN, candesartan; POM, pomegranate.

At baseline (day-1), all rats exhibited normal performance, crossing the beam within 4 to 5 seconds, with no differences between the untreated ischemic animals and those treated with CN alone or CN + POM. following surgery, the untreated ischemic group displayed a significant increase in crossing latency compared with the sham-operated control group (*P* < 0.01 on day +1), which persisted from day +3 to +7 (P < 0.05). In contrast, rats treated with CN or CN + POM exhibited significantly shorter crossing times compared with untreated animals at all post-surgical time points, with no differences from sham controls, indicating complete restoration of motor coordination. There were no differences between CN monotherapy and combined CN + POM treatment, suggesting equivalent efficacy in normalizing beam-walking performance.

### 3.4. Modified sticky tape (MST) test

The MST test evaluated sensorimotor asymmetry and somatosensory-motor integration, reflecting unilateral deficits caused by RCCA ligation-induced cerebral ischemia (**Figure 3**).

**Figure 3.**
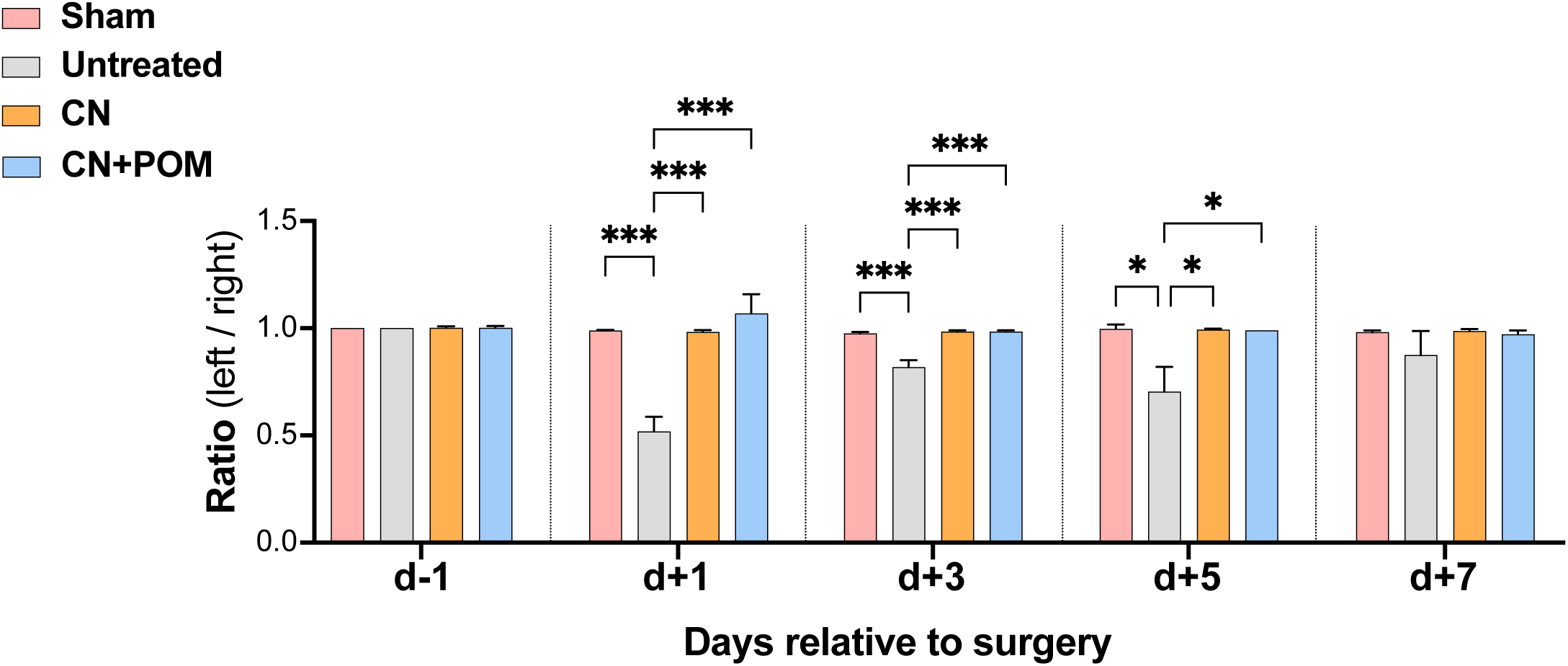
Modified Sticky Tape (MST) test. Sensorimotor asymmetry was evaluated using the modified sticky tape test at baseline (day −1) and on post-operative days +1, +3, +5, and +7. Results are expressed as the left/right tape removal time ratio. Data are presented as mean ± SEM. Statistical comparisons were performed independently at each time point using one-way ANOVA followed by Tukey’s post hoc test (*P < 0.05, ***P < 0.001). *Group and sample size*: Sham (n = 6); Untreated ischemic (n = 6); CN-treated ischemic (n = 5); CN+POM-treated ischemic (n = 6). *Abbreviation*: CN, candesartan; POM, pomegranate.

At baseline (day-1), the left/right removal time ratio was approximately 1 in all groups, indicating symmetrical sensorimotor function (**Figure 3**). Post-surgery, the untreated ischemic group showed a marked reduction in this ratio (∼0.5 on day +1), reflecting a significant unilateral sensorimotor deficit. In contrast, the sham group and animals treated with CN or CN + POM maintained a ratio close to 1, significantly different from untreated ischemic rats (P < 0.001). This difference decreased over time, with the untreated ischemic group gradually approaching a ratio of 1 by day +7, indicating spontaneous recovery.

Overall, treatment with CN alone or in combination with POM prevented the transient sensorimotor asymmetry induced by RCCA ligation.

### 3.5. Novel Object Recognition (NOR) test

The NOR test, performed on post-operative day +8, assessed short-term recognition memory, based on rodents’ spontaneous preference for exploring a novel (N) object over a familiar (F) one (**Figure 4**).

**Figure 4.**
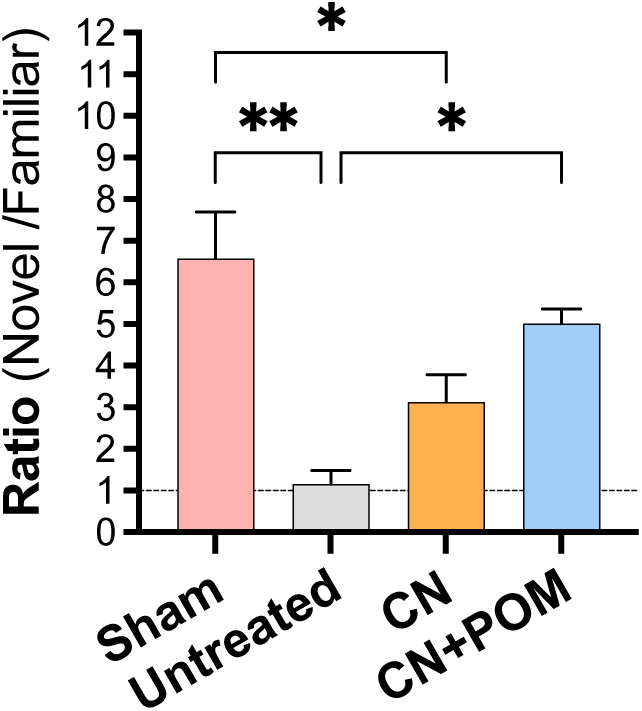
Novel Object Recognition (NOR) test. Short-term recognition memory was assessed on post-operative day +8 using the Novel Object Recognition test. Data are expressed as the recognition ratio (novel/familiar object exploration time). Values are presented as mean ± SEM. Statistical analysis was performed using one-way ANOVA followed by Tukey’s post hoc test (*P < 0.05, **P < 0.01). *Group and sample size*: Sham (n = 6); Untreated ischemic (n = 4); CN-treated ischemic (n = 4); CN+POM-treated ischemic (n = 4). *Abbreviation*: CN, candesartan; POM, pomegranate.

The untreated ischemic group displayed a significantly lower recognition ratio (N/F = 1.2 ± 0.3) compared with sham-operated controls (N/F = 6.6 ± 1.1; P < 0.01), indicating impaired recognition memory. Rats treated with the combined CN + POM therapy showed a significant increase in recognition ratio (N/F = 5.0 ± 0.3; P < 0.05 vs. untreated), reflecting partial restoration of short-term memory.

In contrast, CN monotherapy resulted in a modest, non-significant, increase in recognition ratio (N/F = 3.1 ± 0.7) compared with untreated ischemic rats, indicating lower efficacy than the combined therapy.

### 3.6. Morris water maze (MWM) test

The MWM test assessed spatial learning and memory, based on the ability to locate a hidden platform in a water pool (**Figure 5**).

**Figure 5.**
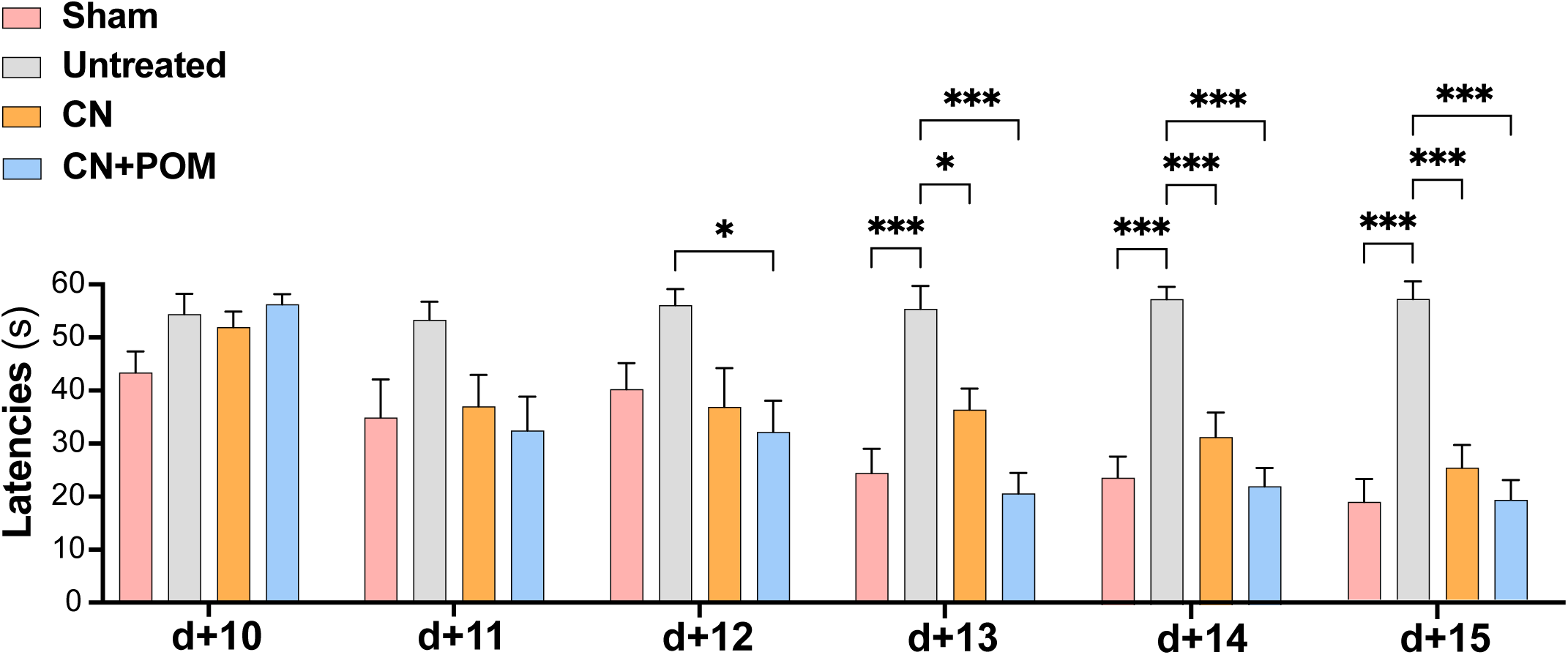
Morris Water Maze (MWM) test. Spatial learning and memory were evaluated using the Morris Water Maze test from post-operative days +10 to +15. Latency to locate the hidden platform was recorded daily. Data are presented as mean ± SEM. Statistical comparisons were performed independently at each time point using one-way ANOVA followed by Tukey’s post hoc test (*P < 0.05, ***P < 0.001). *Group and sample size*: Sham (n = 6); Untreated ischemic (n = 6); CN-treated ischemic (n = 5); CN+POM-treated ischemic (n = 6). *Abbreviation*: CN, candesartan; POM, pomegranate.

On post-operative days +10 and +11, all groups required similar latency times to locate the platform, with no significant differences observed. This indicates that spatial learning deficits were not yet apparent, likely reflecting acute post-operative recovery rather than cognitive impairment, as evidenced by the sham-operated control group showing a gradual decrease in latency from days +12 to +15.

From day +13 to +15, the untreated ischemic group exhibited significantly higher latency times compared with sham controls (e.g., 57.7 ± 2.3 s vs. 24.0 ± 4.0 s at day +14; P < 0.001), establishing the time window for evaluating treatment efficacy.

Within this timeframe, both CN and CN + POM treatments significantly reduced latency times compared with the untreated group, reaching levels similar to sham controls (e.g., 31.7 ± 4.6 s and 22.4 ± 3.5 s at day +14 for CN and CN + POM, respectively). Notably, the combined CN + POM treatment tended to produce a more pronounced effect than CN alone, with a significant reduction in latency detectable as early as day +12 (P < 0.05 *versus* untreated group), while CN monotherapy did not differ significantly from the untreated group at this earlier time point.

## 4. Discussion

Stroke remains a highly prevalent condition worldwide [21] and is often associated with hypoxic or ischemic events that reduced oxygen supply to brain tissue. In this study, we used a rat model of cerebral ischemia induced by ligation of the RCCA. Animals were assigned to four groups: sham control, untreated ischemic, ischemic treated with CN, and ischemic treated with CN combined with POM (CN + POM). Functional outcomes were assessed post-surgery using a battery of behavioral tests, including Beam Balance (BB), Beam Walking (BW), Modified Sticky Tape (MST), Novel Object Recognition (NOR), and Morris Water Maze (MWM).

RCCA ligation produced contralateral sensorimotor deficits, primarily affecting the left side of the body. Both CN and CN + POM treatments fully restored sensorimotor function to sham-control levels, as evidenced by BB, BW and MST performance. In the MST test, the left forelimb (contralateral to the ischemic right hemisphere) was substantially impaired in untreated ischemic rats, requiring roughly twice as long to remove the adhesive tape as the right forelimb, resulting in a left/right ration of ∼0.5. In contract, sham, CN, and CN + POM groups maintained a ratio close to 1, reflecting normal, symmetrical sensorimotor function. These observations indicate that RCCA ligation causes unilateral deficits due to right-hemisphere ischemia, and that CN alone is sufficient to restore contralateral sensorimotor function.

Cognitive deficits were assessed using NOR and MWM tests, which evaluate short-term recognition memory and spatial learning, respectively. RCCA ligation significantly impaired both domains, as indicated by a reduced novel/familiar object recognition ratio and prolonged platform-finding latency in untreated ischemic rats. Importantly, CN + POM treatment produced greater improvements in cognitive performance than CN alone, with earlier and more pronounced recovery in the MWM, whereas in the NOR, only the combined treatment reached significance for improved recovery, and CN alone showed a non-significant trend versus untreated rats.

These findings suggest that the antioxidant and anti-inflammatory properties of POM synergize with CN to provide enhanced protection specifically for cognitive functions, whereas POM does not further enhance sensorimotor recovery provided by CN alone. This highlights distinct mechanisms underlying sensorimotor versus memory recovery following cerebral ischemia.

Previous studies support these observations. Transient global ischemia selectively degenerates hippocampal CA1 neurons 14 days post-ischemia, correlating with severe learning and memory deficits [22]. CN treatment has been shown to improve spatial learning and memory, reduces hippocampal apoptosis, decreases lesion volume, mitigates microglial activation, and preserves functional behavior following brain injury [23–25]. The blockade of angiotensin II type 1 receptors by CN also limits superoxide production and subsequent neuronal damage, with approximately 30% of hippocampal CA1 neurons surviving post-ischemia [22]

The additional neuroprotective effect of POM is likely mediated by its polyphenolic antioxidants, which reduce oxidative stress and inflammation. Lebanese pomegranates cultivated at ∼400 m altitude in the Nabatieh region are particularly rich in polyphenolic compounds compared with those from higher altitudes (700 m in Syria, 600 m in Lebanon-Hasbayah), which may account for their neuroprotective potential. Polyphenolic compounds exhibit neuroprotective activity against oxidative stress. Studies have shown that resveratrol, a polyphenolic compound, protects against spinal cord ischemia-induced damage in rats by eliminating free radicals, exerting antioxidant effects, and blocking the iNOS/p38MAPK signaling pathway [26]. Ischemic events due to oxygenated free radicals decrease motor performance and coordination. The increased synthesis of neurosteroids (antioxidant compounds found in pomegranate) due to the prevention of reactive oxygen species generation may improve motor performance [27]. Additionally, DPPH and phytochemical analyses further confirm the strong antioxidant activity of POM, which can enhance the efficacy of CN [28]. Similar synergistic effects have been observed with CN combined with other antioxidants, such as curcumin, supporting the concept that antioxidant therapy can augment AT1R blockade to improve post-ischemic recovery [1].

Overall, our results highlight a differential contribution of CN and POM: CN effectively restores sensorimotor function, while POM provides synergistic and specific protection against cognitive deficits, emphasizing complementary mechanisms underlying motor versus memory recovery after ischemic injury.

Several limitations should be noted. First, the study relied exclusively on behavioral assessments, without histological (e.g., infarct volume) or biochemical analyses (e.g., oxidative stress or inflammatory markers) to elucidate underlying cellular mechanisms. Second, the unilateral RCCA ligation model in rats may produce variable ischemic severity due to collateral circulation, limiting extrapolation to human stroke. Third, we did not assess the effect of POM alone, which would be necessary to determine its independent contribution to neuroprotection. Finally, translation to clinical practice requires further investigation of pharmacokinetics, long-term safety, and dose-response interactions in humans.

## 5. Conclusions

This study demonstrates that combined treatment with CN and pomegranate juice provides superior neuroprotection compared to CN monotherapy in a rat model of cerebral ischemia. While RCCA ligation induces profound sensorimotor deficits and cognitive impairment, CN alone restores sensorimotor function, and the addition of POM enhances recovery of memory and spatial learning. These findings highlight the synergistic and complementary effects of AT1R blockade by CN and the antioxidant properties of POM, suggesting that this dual-treatment approach may be a promising strategy for mitigating post-ischemic brain injury and improving functional outcomes.

## Abbreviations

AT1R: Angiotensin II type 1 receptor
CN: Candesartan
POM: Pomegranate
RCCA: Right common carotid artery
BB: Beam balance
BW: Beam walking
MST: Modified sticky-tape
NOR: Novel object recognition
MWM: Morice water maze
RAS: Renin-angiotensin system
ROS: Reactive oxygen species.

## Availability of Data and Materials

The data presented in this study are available on request from the corresponding authors.

## Author Contributions

M.N, conceptualization; AH, methodology; FR and M.N data analysis; L.O.M and M.N, validation; F.R, A.H and M.N, formal analysis; W.J, investigation; R.A and M.N, resources; F.R and M.N, data curation; R.A and M.N, writing original draft preparation; F.R, Z.E, Z.A.B, L.O.M and M.N, writing review and editing; A.H and MN, supervision; W.J and M.N, project administration; AH, L.O.M and M.N, funding acquisition. All authors have read and agreed to the published version of the manuscript.

## Ethics Approval and Consent to Participate

Animal experiments were conducted humanely in accordance with the Regulations for Animal Experiments of Islamic University of Lebanon which are in accordance with the revised Animals Act 1986 in the UK and Directive 2010/63/EU in Europe). The study was approved by the research ethics committee at the Islamic University of Lebanon (reference number: IUL-EC-25-A005)

## Acknowledgment

This work was supported by the Lebanese university and by the French Foreign Ministry through the Hubert Curien Project - Cèdre N°42232RE. The authors are also thankful to the High Council for Scientific Research & Publication (HCSRP), Islamic University of Lebanon.

