## Supplementary material for "Neuroprotective Effect of Combined Pomegranate and Candesartan Therapy Against Chronic Cerebral Ischemia in Rats": Table S1 and S2

**Table S1. Screening methods for the detection of primary and secondary metabolites in pomegranate juice and seeds.**

| Plant constituents | Reagents added | Color |
|---|---|---|
| Phenolic acids | $FeCl_3$ (1%) + $K_3(Fe(CN)_6)$ (1%) | Greenish Blue |
| Terpenoids | Chloroform + concentrated sulfuric acid | Reddish brown in the surface |
| Flavonoids | KOH (potassium hydroxide 50%) | Yellow |
| Quinones | HCl concentrated | Precipitate or yellow |
| Alkaloids | Dragendroff reagent | Reddish Orange precipitate/turbidity |
| Tannins | $FeCl_2$ (1%) | Blue |
| Resins | Acetone + water + agitation | Turbidity |
| Saponins | Vigorous shaking | Layer of foam |
| Reducing sugar | Water + fehlings (A+B) + boil | Brick-red precipitate |
| Anthraquinones | HCl (10%) + boil | Precipitate |
| Proteins and Amino Acids | Ninhydrin (0.25%) + boil | Blue |
| Phlabotannins | HCl (1%) + boil 5 min + cooling | Red precipitate |
| Flavanones | $H_2SO_4$ concentrated | Purple red |
| Diterpenes | Copper sulfate | Green |
| Sterols and Steroids | Chloroform + $H_2SO_4$ concentrated | Red color (upper layer) and greenish yellow fluorescence (acidic layer) |
| Anthocyanins | NaOH (10%) | Blue |
| Lignins | Safranin | Pink |
| Cardiac glycosides | Acetic acid glacial + $FeCl_3$ (5%) + concentrated $H_2SO_4$ | Purple ring + brown ring + green ring |
| Fixed oils and fatty acids | Spot test | Oil spot |

Each plant constituent was identified using specific reagents and characterized by its expected colorimetric or physical reaction (color change, precipitate, fluorescence, or turbidity).

**Table S2. Chemical composition of pomegranate juice and seeds.**

| Bioactive molecule | Juice | Seeds |
|---|---|---|
| Reducing sugar | - | + |
| Anthraquinones | - | - |
| Proteins and amino acids | - | - |
| Phlabotannins | - | - |
| Alkaloids | + | + |
| Tanins | + | - |
| Resins | - | + |
| Terpenoids | + | + |
| Flavonoids | + | + |
| Quinones | - | + |
| Sterols et steroids | + | + |
| Diterpenes | - | - |
| Anthocyanins | + | - |
| Flavanones | + | + |
| Lignins | + | + |
| Cardiac glycosides | - | + |
| Saponins | - | - |
| Phenols | + | + |
| Fixed oils and fatty acids | - | - |

"+": compound detected/present; "−": compound not detected/absent. Phytochemical classes were assessed by standard colorimetric and qualitative screening methods. Results reflect the presence or absence of each molecule class rather than quantitative levels.
